# Epigenetic clock and methylation study of oocytes from a bovine model of reproductive aging

**DOI:** 10.1101/2020.09.10.290056

**Authors:** Paweł Kordowitzki, Amin Haghani, Joseph A. Zoller, Caesar Z. Li, Ken Raj, Matthew L. Spangler, Steve Horvath

## Abstract

Cattle are an attractive animal model of fertility in women due to their high degree of similarity relative to follicle selection, embryo cleavage, blastocyst formation, and gestation length. To facilitate future studies of the epigenetic underpinnings of aging effects in the female reproductive axis, several DNA methylation-based biomarkers of aging (epigenetic clocks) for bovine oocytes are presented. One such clock was germane to only oocytes, while a dual-tissue clock was highly predictive of age in both oocytes and blood. Dual species clocks that apply to both humans and cattle were also developed and evaluated. These epigenetic clocks can be used to accurately estimate the chronological age of the oocyte donor. Both epigenetic clock studies and epigenome wide association studies revealed that blood and oocytes differ substantially with respect aging and the underlying epigenetic signatures that potentially influence the aging process. The rate of epigenetic aging was found to be slower in oocytes compared to blood, however, oocytes appeared to begin at an older epigenetic age. The epigenetic clocks for oocytes are expected to address questions in the field of reproductive aging, including the central question: how to slow aging of oocytes.

## INTRODUCTION

The mammalian female reproductive axis is the first to fail in aging, however, the molecular mechanisms underpinning this failure are largely unknown, particularly in oocytes [1-3]. Fertility in women begins to decline significantly by their mid-30s and pregnancies in women of advanced age lead to higher rates of miscarriage and/or aneuploid offspring [4]. Despite these risks, women and couples often postpone pregnancy to a more convenient time which has led to a decline in birthrate among most industrialized societies [1,5]. This decline in fertility can be explained by the age-related decline in oocyte quality which manifests itself by chromosomal abnormalities, spindle defects, mitochondrial dysfunction, and epigenetic modifications [4, 6-11]. Due to ethical restrictions for research on human oocytes, bovine oocytes are a common and attractive model of oocyte developmental competence and reproduction. Interest in bovine pre-implantation embryology and bovine/ruminant *in vitro* models in the field of human reproduction has increased [12, 13].

Oocytes of donors with increased maternal age also show aberrant global DNA methylation levels [14]. Previous work has shown that highly accurate estimators of chronological age (epigenetic clocks) that apply to most tissues and cell types can be developed using DNA methylation levels at individual loci [15, 16]. The current study addresses several key questions surrounding the development of an epigenetic clock of oocytes. Since oocytes can be readily collected in cattle, DNA from both cattle and humans were used to address the following aims: 1) to test whether an epigenetic clock that applies to blood also applies to oocytes, 2) to build an epigenetic clock for bovine oocytes, 3) to build epigenetic clocks that apply to both cattle and humans to facilitate translation of research findings, 4) to evaluate whether epigenetic age acceleration detected in blood is also manifested in oocytes, and 5) to ascertain the association between age and methylation levels of individual CpGs in oocytes.

## RESULTS

DNA methylation data were generated from DNA samples (n=357) from oocyte donors (n=80) and blood donors (n=277) of cattle (*Bos taurus*). Animals were between 0.5 years and 13.3 years of age at the time of tissue sampling. Unsupervised hierarchical clustering revealed that oocytes and blood methylation profiles form distinct clusters (Supplementary Figure 3). Using penalized regression, several highly accurate epigenetic clocks were developed for cattle blood, cattle oocytes, and both tissues combined (Figure 1A-C). These epigenetic clocks differ with regards to their operational parameters; tissue, species and measure of age.

**Figure 1:**
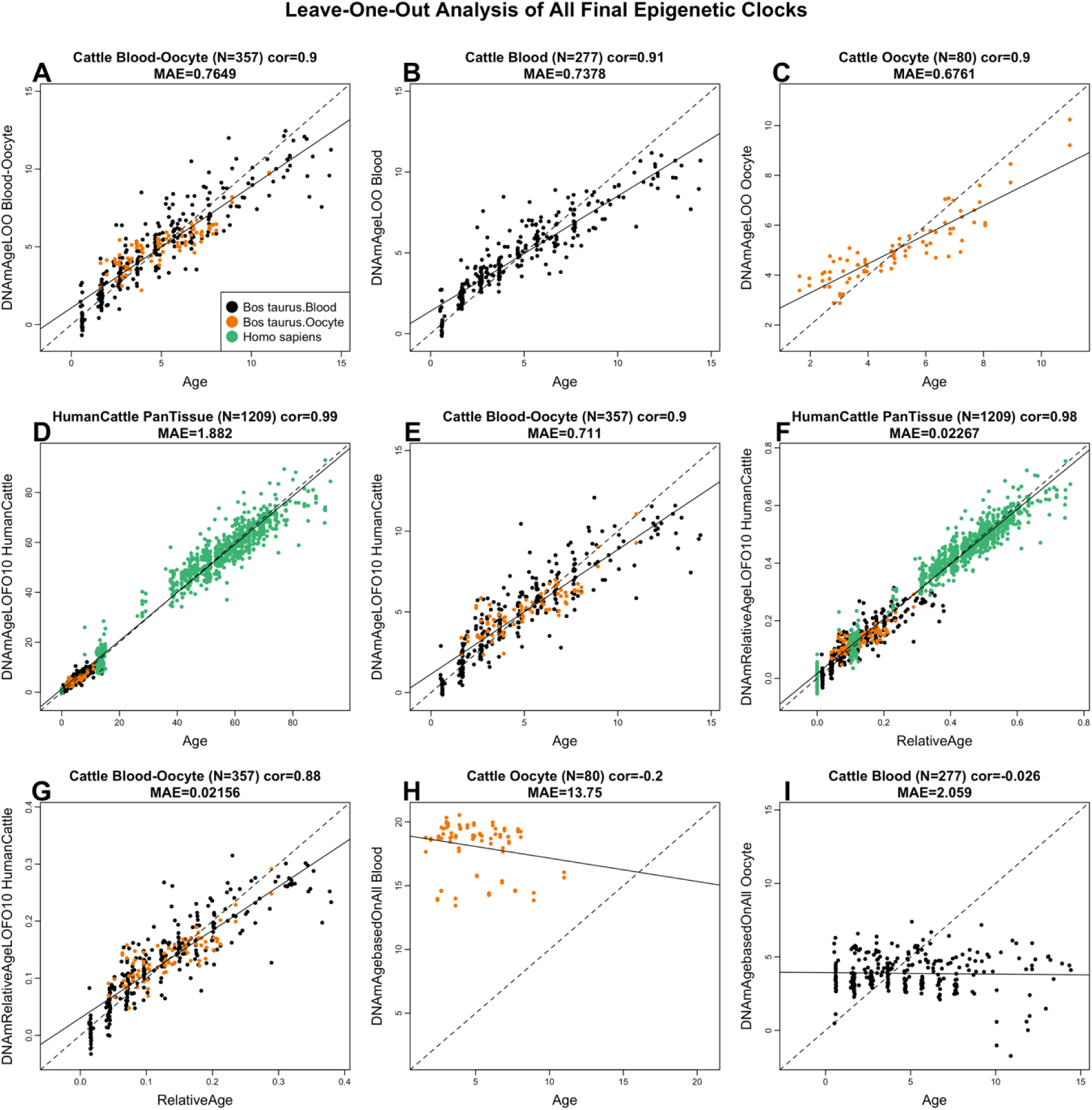
Cross-validation study of epigenetic clocks for cattle and humans. A) Blood-oocyte clock for cattle applied to blood and oocytes, B) Cattle clock for blood, C) Cattle clock for oocytes. Chronological age (x-axis, in units of years) versus the leave-one sample-out (LOO) estimate of DNA methylation age (y-axis, in units of years). Dots are colored by tissue type (orange=oocytes, black=blood). D,E) Ten-fold cross validation analysis of the human-cattle clock for (chronological) age applied D) to samples from both species and to E) to cattle only. F,G) Human-cattle clock estimate of relative age defined as the ratio of chronological age to the maximum lifespan of the respective species. Ten-fold cross-validation estimates of age (y-axis, in years) in D,F) Human (green) and cattle (orange) samples and E, G) cattle samples only (colored by tissue type). Each panel reports the sample size, correlation coefficient, median absolute error (MAE). H) Cattle blood clock applied to oocytes from cattle. I) Cattle oocyte clock applied to blood samples from cattle.

To develop dual species clocks, DNA methylation profiles from human samples (n=852) were added to the training set. Methylation data for both bovine and human tissues were generated from the HorvathMammalMethylChip40 array that consists of approximately 36000 CpGs embedded in DNA sequences that are highly conserved within the mammalian class. As would be expected, a high percentage (45%) of conserved genes and regions between cattle and humans was found (Supplementary Figure 1). The mammalian array coverage of the cattle genome (*Bos taurus*) ARS-UCD1.2 based on genome alignments with the human Hg38 genome is reported herein (Supplementary Figure 1). In total, 34331 out of 37540 probes mapped to the cattle genome and approximately 64% of these mapped to the same genes in cattle and humans (Supplementary Figure 1). This high degree of conservation suggests that building dual species clocks that apply to both humans and cattle is indeed possible. The two human-cattle clocks that were derived, used the same set of CpGs and the same prediction equation to estimate age in both species. The dual species clocks presented herein can be distinguished by the measure of age; whereby one operates with chronological age in units of years, while the other employs relative age, which is the ratio of chronological age to the maximum lifespan of the species, and is expressed as values between 0 and 1. The maximum ages of cattle and humans are 38 years and 122.5 years, respectively. The high accuracy of the human-cattle clock for relative age (Figure 1F,G) shows that this clock facilitates a biologically meaningful comparison between species with different lifespans (cattle and human). The human-cattle clock generated a correlation of r=0.99 between chronological age and epigenetic age when both species were analyzed together (Figure 1D), but the correlation was slightly lower when the analysis was restricted to bovine oocytes and blood samples (r=0.90, Figure 1E). The results of the human-cattle clock for relative age show a comparably high correlation regardless of whether the analysis was performed with samples from both species (r=0.98, Figure 1F) or only with bovine samples (r=0.88, Figure 1G). By construction, the multi-tissue clock for blood and oocytes applies to both sources of DNA (Figure 1A). Interestingly, the cattle clock for blood does not apply to oocytes (Figure 1H) and, conversely, the cattle clock for oocytes does not apply to cattle blood (Figure 1G).

As a subset of data were generated from blood and oocytes from the same animal, epigenetic ages between these two isogenic tissues were directly compared. To avoid the confounding effect of chronological age, estimated DNAm age acceleration was used as the feature of comparison for both tissues. None of the epigenetic clocks resulted in a significant positive correlation between epigenetic age acceleration in blood and that of oocytes from the same cattle (Figure 2).

**Figure 2.**
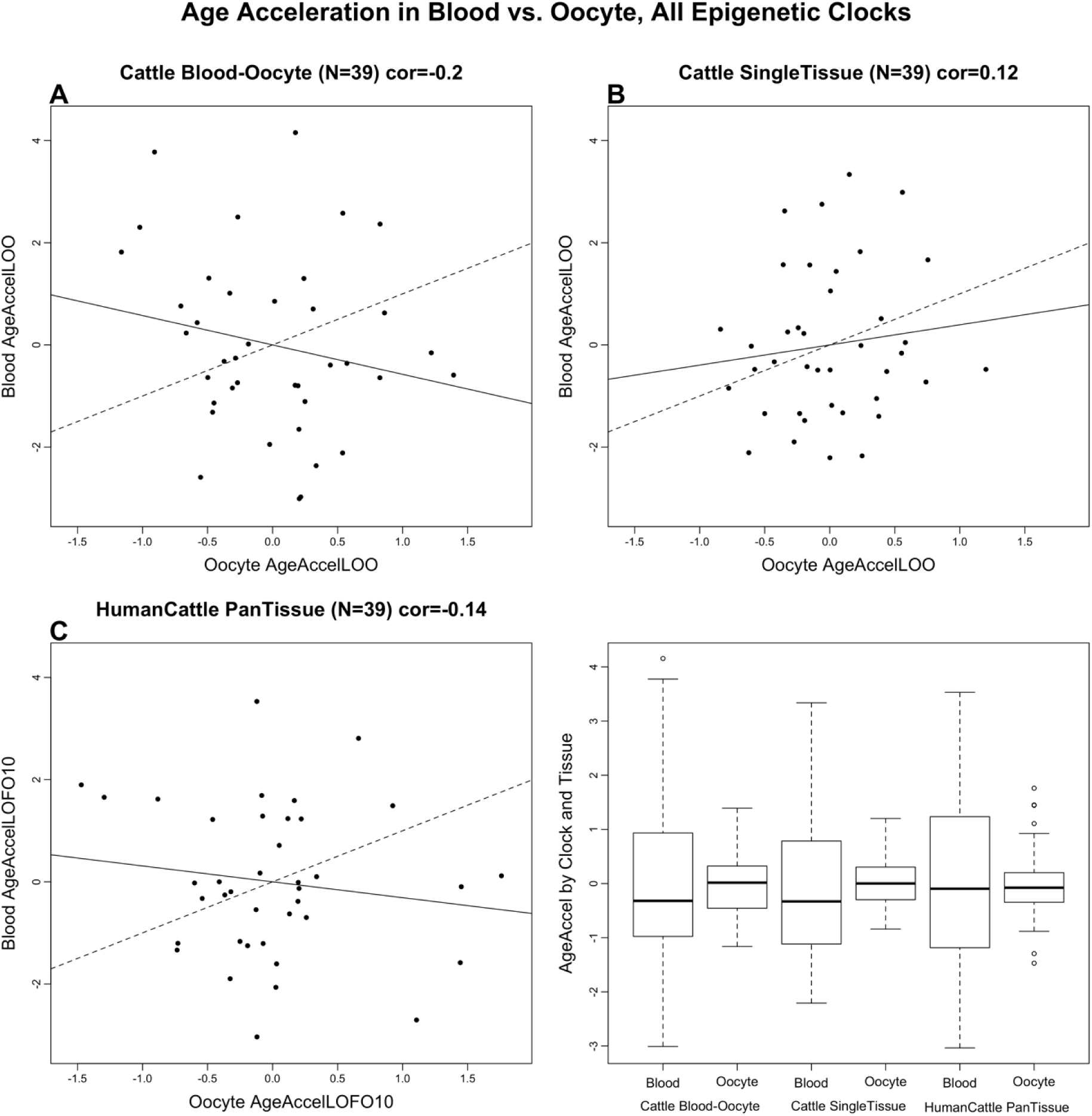
Epigenetic age acceleration in cattle blood is not correlated with that in oocytes. A-B) Cross validation estimates of epigenetic age acceleration in blood versus epigenetic age acceleration in oocytes. Dashed line indicates the diagonal line (y=x). The solid line corresponds to the regression line. A) Results for the multi-tissue cattle clock. B) Single tissue clocks for oocytes (x-axis) and blood (y-axis). C) Human-cattle clock of chronological age. D) Boxplots of epigenetic age acceleration in blood or oocytes for different cattle clocks.

The epigenetic clocks for cattle performed poorly when used to predict the age of human blood and skin samples (Supplementary Figure 5). The best performing of the three pure cattle clocks was the multi-tissue clock whose age estimates were only moderately correlated with age in human blood (r=0.38) and skin (r=0.5) samples (Supplementary Figure 5A,D). As expected, the human-cattle clocks led to substantially higher age correlations in human samples (cross validation estimates of r=0.99 in both blood and skin, Supplementary Figure 6).

An epigenome-wide association study (EWAS) on chronological age in bovine blood and oocyte samples was also conducted. The results revealed that blood and oocytes have distinct age-dependent DNA methylation changes. Individual CpGs exhibited highly significant age correlations in blood and oocytes (Figure 3A). Results from the EWAS showed that individual CpGs in cattle blood (Figure 3A), oocytes (Figure 3B) were highly associated with age. Oocytes and blood have distinct sets of age-related CpGs; with oocytes having considerably fewer age-related CpGs (Figure 3, Supplementary Figure 7A,C). Although more than 500 CpGs were hypermethylated and more than 500 were hypomethylated in blood (p < 10^−4^), only 71 and 75 CpGs were hypermethylated and hypomethylated, respectively, in oocytes at the same significance threshold (Figure 3B-C). This striking difference in the number of age related CpGs between tissue types was also observed when restricting the analysis to blood and oocyte samples from the same animals (Supplementary Figure 7). The top DNAm changes are as follows: hypermethylation in *NBEA* intron in blood (Figure 4A), and hypomethylation in *TCF20* downstream region in oocytes (Figure 4B). In the meta-analysis of blood and oocytes, the top DNAm change was also hypermethylation in NBEA intron (Figure 3A, Figure 4C). One of the top DNAm changes in oocytes was near the *DENND1A* gene (Table 1, chromosome 11). Genetic variants in *DENND1A* have been suggested to play a role in susceptibility to polycystic ovary syndrome (*PCOS*), the most common endocrine disease among premenopausal women. PCOS is a complex disorder characterized by infertility, hirsutism, obesity, various menstrual disturbances, and enlarged ovaries studded with atretic (degenerated) follicles [17-20].

**Table 1.** Top 40 negatively and positively age associated CpGs in oocyte.

| CGid | SYMBOL | Region | Chromosome | Z score | P | Conservation in human |
| --- | --- | --- | --- | --- | --- | --- |
| cg12593479 | PITX2 | Exon | 6 | 5.32 | 1.06E-07 | different |
| cg04734735 | HOXD8 | Exon | 2 | 5.17 | 2.32E-07 | different |
| cg20627174 | EN1 | Intergenic_downstream | 2 | 5.14 | 2.71E-07 | conserved |
| cg21811143 | EN1 | Intergenic_downstream | 2 | 4.98 | 6.29E-07 | conserved |
| cg09009312 | DENND1A | Intron | 11 | 4.97 | 6.66E-07 | conserved |
| cg03461257 | DENND1A | Intron | 11 | 4.89 | 1.03E-06 | conserved |
| cg21215767 | EN1 | Intergenic_downstream | 2 | 4.62 | 3.83E-06 | conserved |
| cg17892435 | PITX2 | Intron | 6 | 4.62 | 3.93E-06 | conserved |
| cg19688271 | MKI67 | Intergenic_upstream | 26 | 4.53 | 6.01E-06 | different |
| cg14467682 | EN1 | Intergenic_downstream | 2 | 4.52 | 6.28E-06 | conserved |
| cg17245416 | NOG | Intergenic_downstream | 19 | 4.49 | 6.98E-06 | conserved |
| cg09671306 | SYNGAP1 | Intron | 23 | 4.48 | 7.64E-06 | conserved |
| cg22222486 | FOXO6 | Intergenic_upstream | 3 | 4.47 | 7.97E-06 | conserved |
| cg12911542 | HOXA6 | Promoter | 4 | 4.45 | 8.62E-06 | different |
| cg11375458 | HOXD8 | Intergenic_downstream | 2 | 4.44 | 9.13E-06 | different |
| cg25379954 | PROX1 | Exon | 16 | 4.42 | 9.97E-06 | conserved |
| cg24949381 | DMRT1 | Intron | 8 | 4.39 | 1.11E-05 | conserved |
| cg25528941 | MAFF | Exon | 5 | 4.39 | 1.16E-05 | conserved |
| cg11040386 | IFFO2 | Exon | 2 | 4.38 | 1.17E-05 | conserved |
| cg25162386 | MAFF | Exon | 5 | 4.37 | 1.24E-05 | conserved |
| cg26899365 | ARID1B | Intron | 9 | -4.50 | 6.93E-06 | conserved |
| cg08676154 | ERI3 | Intron | 3 | -4.50 | 6.68E-06 | conserved |
| cg15419333 | ZFHX3 | Intergenic_downstream | 18 | -4.50 | 6.68E-06 | different |
| cg22825448 | DMBX1 | Promoter | 3 | -4.64 | 3.44E-06 | conserved |
| cg20369157 | CMC1 | Intergenic_upstream | 22 | -4.70 | 2.64E-06 | conserved |
| cg06426413 | LMO1 | Intergenic_upstream | 15 | -4.70 | 2.63E-06 | different |
| cg24981538 | ARHGAP15 | Intron | 2 | -4.72 | 2.40E-06 | conserved |
| cg11393097 | MAPKAP1 | Intron | 11 | -4.74 | 2.11E-06 | conserved |
| cg06894182 | POU4F3 | Intergenic_downstream | 7 | -4.75 | 2.04E-06 | conserved |
| cg22968128 | ETV5 | Intron | 1 | -4.77 | 1.86E-06 | conserved |
| cg21621926 | URI1 | Intergenic_downstream | 18 | -4.78 | 1.78E-06 | conserved |
| cg22158106 | MAPKAP1 | Intron | 11 | -4.78 | 1.71E-06 | conserved |
| cg12544505.2 | TCF20 | Intergenic_upstream | 5 | -4.79 | 1.68E-06 | conserved |
| cg11133252 | TCF20 | Intergenic_upstream | 5 | -4.80 | 1.62E-06 | conserved |
| cg27135106 | URI1 | Intergenic_downstream | 18 | -4.85 | 1.26E-06 | conserved |
| cg20814160 | FAM193B | Exon | 7 | -4.93 | 8.10E-07 | conserved |
| cg02016833 | PMFBP1 | Intergenic_upstream | 18 | -5.15 | 2.66E-07 | different |
| cg26481779 | MAPKAP1 | Intron | 11 | -5.28 | 1.29E-07 | conserved |
| cg00472926 | VRK1 | Intergenic_downstream | 21 | -5.28 | 1.26E-07 | different |

**Figure 3.**
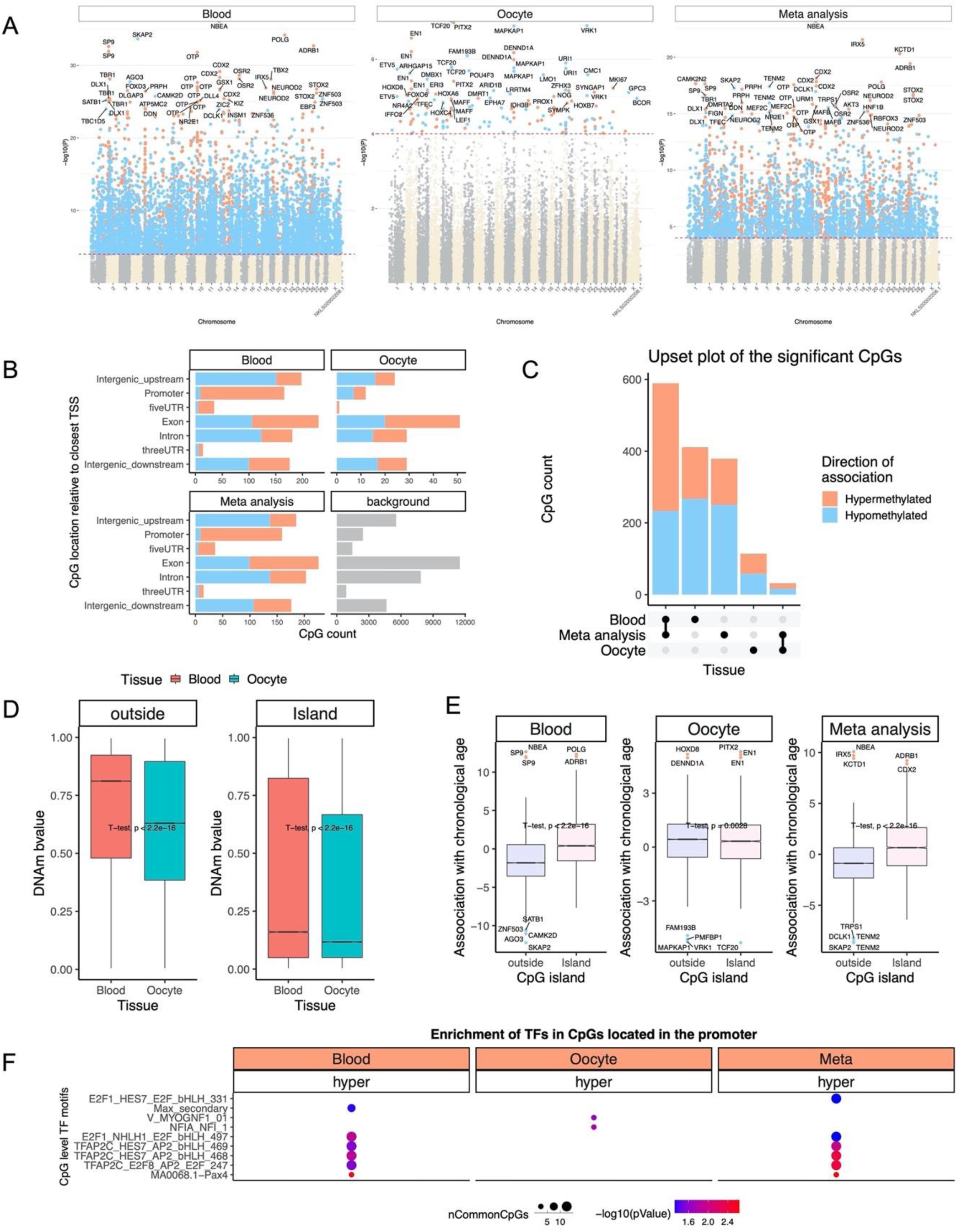
Blood and oocyte have distinct age-dependent DNA methylation changes. A) Manhattan plots of the EWAS of chronological age. The coordinates are estimated based on the alignment of Mammalian array probes to the Bos_taurus.ARS-UCD1.2 genome assembly. The red dotted line corresponds to a significance threshold of p < 10^−4^. Individual CpGs are colored in red or blue if they gain or lose methylation with age. The 30 most significant CpGs are labeled by neighboring genes. B) Location of top CpGs in each tissue is relative to the closest transcriptional start site. Top CpGs were selected (p < 10^−4^) and further filtering based on z score of association with chronological age for a maximum 500 in each direction (positive and negative). The number of selected CpGs: blood, 1000; oocyte, 146; meta-analysis, 1000. The grey color in the last panel represents the location of 34331 mammalian BeadChip array probes mapped to Bos_taurus.ARS-UCD1.2 genome. C) Upset plot representing the overlap of aging-associated CpGs based on meta-analysis or individual tissues. Neighboring genes of the overlapping CpGs were labeled in the figure. D) CpG island analysis restricted to animals for whom both blood and oocyte samples were available. Oocytes have generally lower DNAm levels both within and outside of CpG islands than blood. The mean difference was examined by t-test. E) Box plot of age correlation test Z statistics versus CpG island status in blood and oocytes.. F) Transcriptional motif enrichment for the top CpGs in the promoter and 5’UTR of the neighboring genes. The motifs were predicted using the MEME motif discovery algorithm, and the enrichment was tested using a hypergeometric test.

**Figure 4.**
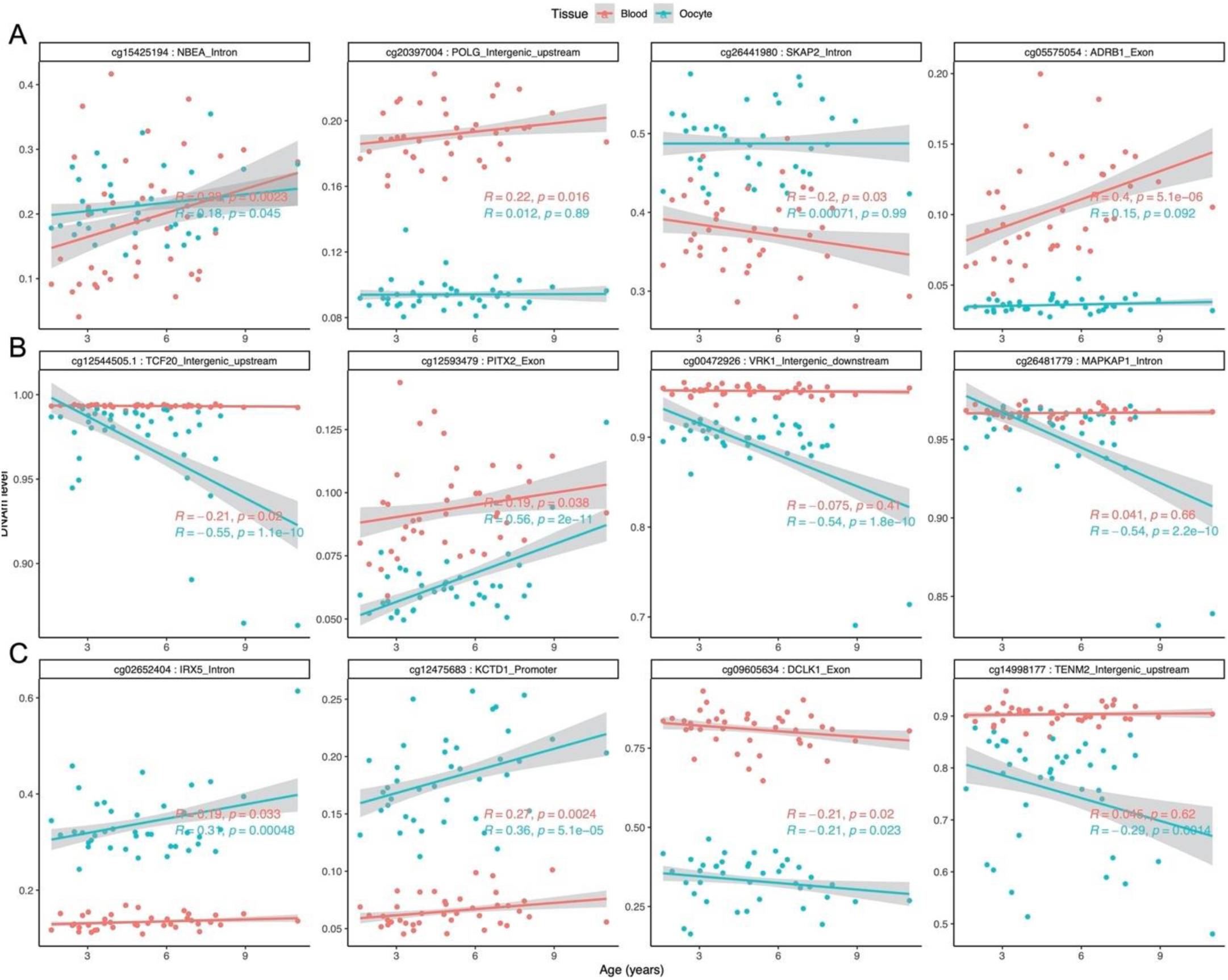
Scatter plots of the most significnatly age-associated CpGs in cattle blood and oocytes. A) top four DNAm age changes in aging blood. cg15425194 and cg05575054 were also identified by meta-analysis. B) top DNAm age changes in aging oocytes. C) Additional DNAm changes identified by meta-analysis.

Across all CpGs, age effects in cattle blood exhibited a weak negative correlation (r=-0.11) with those in oocyte (Supplementary Figure 4). Although a conservative p-value threshold of 10^−4^ was used to define age related CpGs, findings are qualitatively the same for other significance thresholds. Striking differences between blood and oocytes were observed among the significant age related CpGs. First, age-related CpGs in gene promoters exhibited positive age correlations in blood but not in oocytes (odds ratio=23.3, hypergeometric test p=7.2⨯10^−7^, Figure 3B). Second, none of the age-related CpGs were shared between blood and oocytes (Figure 3C). For CpGs outside of CpG islands, the mean methylation level in oocytes was substantially lower than that of blood (Figure 3D). The difference was less pronounced for island CpGs. (Figure 3D). In blood, island CpGs exhibited substantially higher positive age associations than CpGs outside of islands (Figure 3E). This well known pattern [21] could *not* be observed in oocytes. Rather, non-island CpGs in oocytes appear to be refractive to demethylation with age (Figure 3E), which may reflect that their ground-state of methylation was already low and could not be further reduced. Enrichment of human tissue-specific epigenome states suggested that hypermethylated CpGs in blood are located near bivalent/poised transcription start sites, flanking bivalent TSS, bivalent enhancers, and repressed polycombs (Supplementary Figure 8). These features were similarly observed in humans [21]. Histone 3 marks for blood hypermethylated CpGs included H3K27me3, H3K4me1, and H3K9me3. Age-related CpGs found in oocytes were not enriched for any epigenetic signatures of tissue type. By contrast, age-related CpGs found in blood were, as expected, enriched for epigenetic signatures of blood (Supplementary Figure 8). Transcription factor binding sites that were differently methylated with age were also distinct between blood and oocytes. The highest scoring transcription factor motifs are hypermethylation in PAX4 and TFAP2C binding regions (Figure 3F). In contrast, the highest score for oocytes were hypermethylation in NFIA and MYOGNF1-binding motifs. Enrichment analysis of genes proximal to the top age-related CpGs in blood and oocyte is shown in Supplementary Figure 2.

## DISCUSSION

Given clear differences in aging effects between oocytes and blood, it was hypothesized that methylation of select CpGs in these two tissues may move in opposing directions with age. To test this hypothesis, a multivariate model was employed to analyse blood and oocyte samples derived from the same animals. A total of 45 CpGs exhibited significant interaction effects between age and sample type at a nominal significance of p <0.001 (Figure 5A). The aging patterns of these CpG sites were classified into 9 types (Figure 5B). Interestingly, CpGs within *DENND1A* intron were among those that changed with age in both oocyte and blood (Figure 5C), but while they became increasingly hypomethylated with age in blood, they were hypermethylated in aging oocytes. The enrichment analysis of this set of CpGs also highlighted differences in Rap GTPase, which plays an essential role in metabolic processes and signal transduction [19]. The Rap1 mutation leads to infertility by impairing germ-Sertoli cell contacts, and higher expression in women with premature ovarian failure has been reported [20].

**Figure 5.**
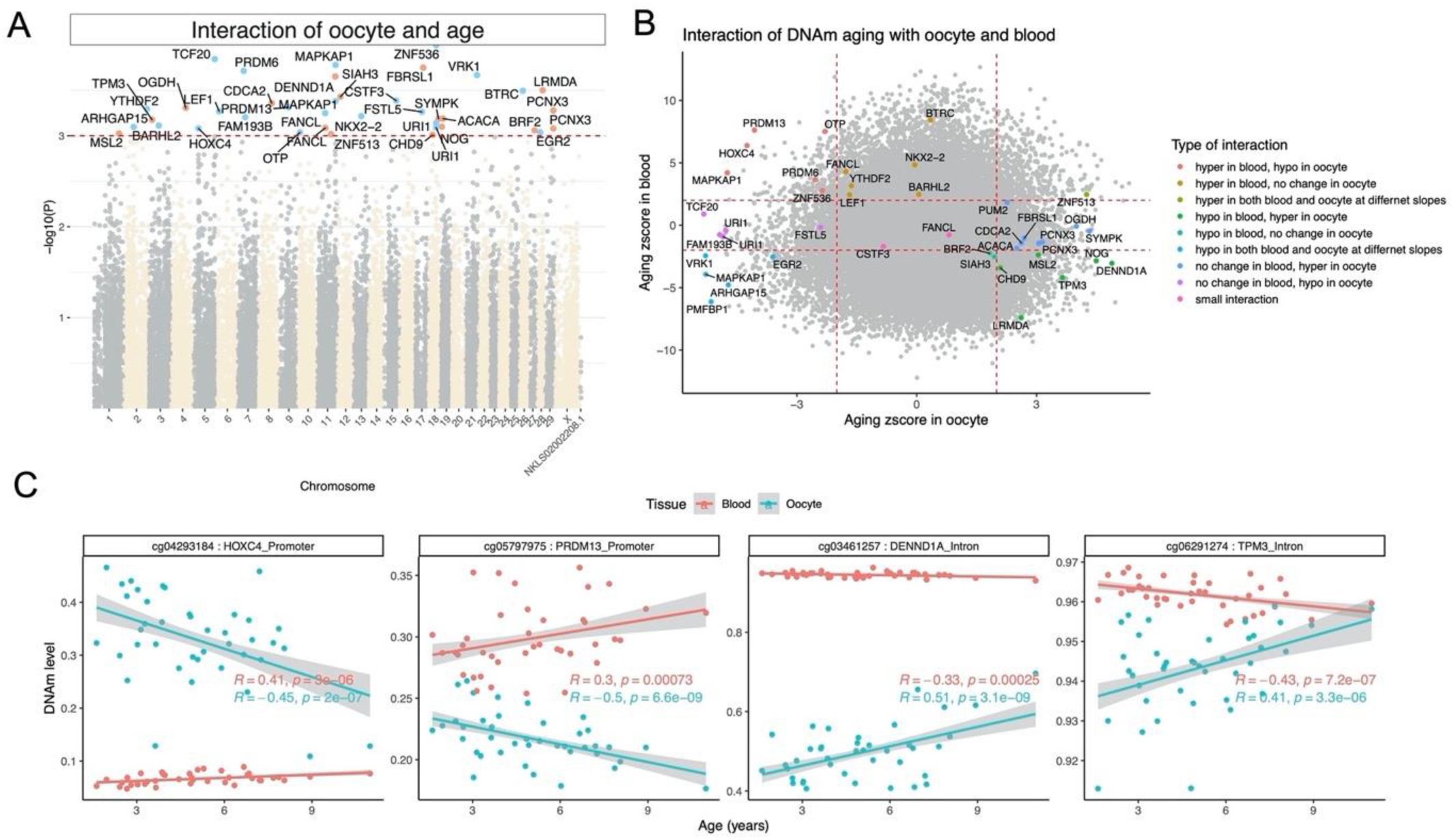
DNAm aging is impacted by sample type (oocyte and blood) at several specific CpGs. Multivariate model analysis for detecting an interaction effect between age and sample type in cows (n=40). Dependent variable: individual CpG. Covariates: age, sample type, and the interaction term age x sample type. A) Manhattan plot of the log (based 10) p value of the interaction effect. 45 CpGs passed the significance threshold p < 10^−3^ (red dotted line). B) Nine different types of interactions are depicted for DNAm between blood and oocyte based on the Z statistics of an age correlation analysis stratified by sample type. C) Scatter plots of selected CpGs with significant interaction effects between age and sample type.

Recent evidence suggests that SIRT1 plays a role in oocyte function in older ages [22]. The mammalian array used in the current study did not cover the *SIRT1* gene in the cattle genome, but CpGs in the 1Mb flanking region of the *SIRT1* gene could be investigated (Supplementary Figure 9). Several CpG probes downstream of *SIRT1* were hypomethylated with age in both blood and oocytes. Interestingly, a probe upstream of *SIRT1* was only hypomethylated with age in oocytes but not in blood.

The stark difference in multiple characteristics of age-related CpGs between blood and oocytes is perhaps not surprising. What is unexpected is the successful development of oocyte-based epigenetic clocks including dual tissue clocks that apply to both blood and oocytes. Methylation of DNA plays a well known role in genomic imprinting, which is one of the most important events in the development and maturation of oocytes. The mammalian array covered 23 CpG probes proximate to seven known parental imprinted genes (DCN, GNAS, PEG10, NAA60, MES, PLAGL1, and SFMBT2) in cattle. None of the observed DNAm aging loci in blood or oocytes overlapped with these 7 imprinted genes. Overall, the potential link between DNAm aging and the imprinting mechanism in oocytes could not be elucidated in this study.

If the oocyte epigenetic clocks turn out to be indicators of the fitness of oocytes, it would suggest the existence of a maternal age-dependent program that specifically alters the DNA methylation state of the oocyte. The endocrine system, which changes with age, could relate to this program. Indeed, age-related hormonal changes are very well-documented, and it is worth noting the recent report on the rejuvenation effects of human growth hormone [23]. Regardless of the actual mechanism, which requires empirical elucidation, the potential importance of age-related DNA methylation changes on oocyte DNA cannot be ignored. The epigenetic clocks presented herein for oocytes are expected to be useful for finding answers to many questions that range from fecundity to evolutionary selection.

## CONCLUSION

The primary objective of the study was to develop and apply epigenetic biomarkers of aging for oocytes from cattle as a model for human oocytes. Although a dual-tissue epigenetic clock was developed that was predictive of age in both blood and oocytes, the fundamental aging properties of these two sample types were found to differ substantially. The rate of epigenetic aging was found to be slower in oocytes compared to blood, however, oocytes appeared to begin at an older epigenetic age. The differences in epigenetic aging effects between oocytes and blood were observed at the level of individual CpGs and cumulatively at the level of single-tissue epigenetic clocks. The epigenetic clocks for oocytes are expected to address questions in the field of reproductive aging, including the central question: how to slow aging of oocytes.

---

**Supplementary Figure 1.**
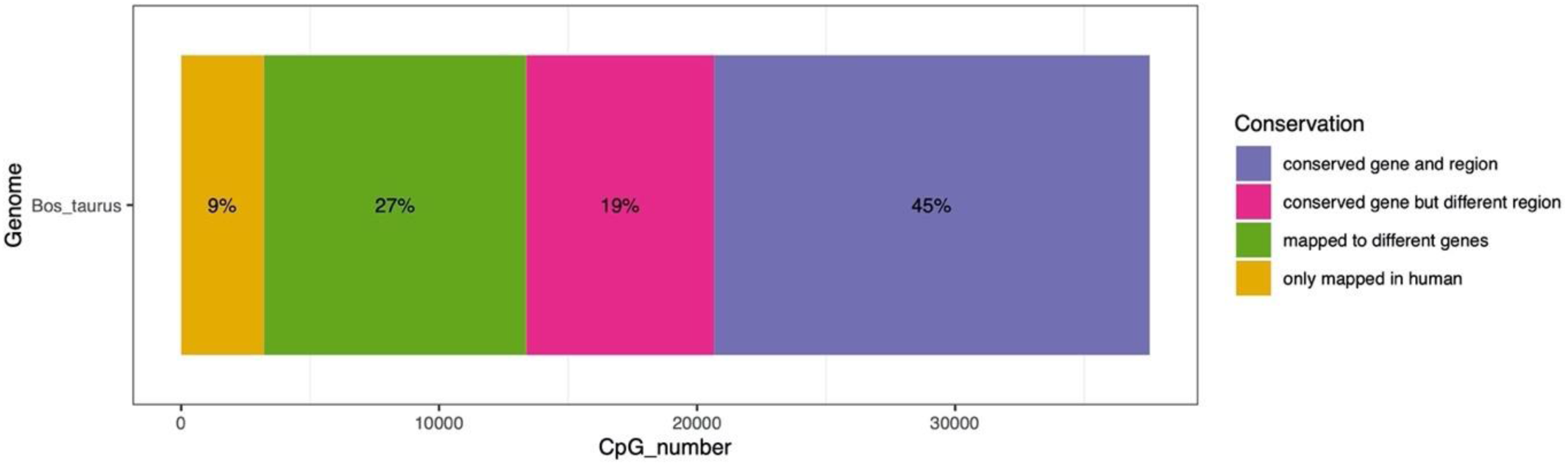
Coverage of mammalian array on Cattle (*Bos taurus*) ARS-UCD1.2 genome. In total, 34331 out of 37540 probes could be mapped to the cattle genome. The stacked bar compares the alignments with the human Hg38 genome. Approximately 64% of the probes were mapped to the same genes between cattle and humans. 17% of the probes that mapped to different genes in these species were located in distal intergenic regions.

**Supplementary Figure 2.**
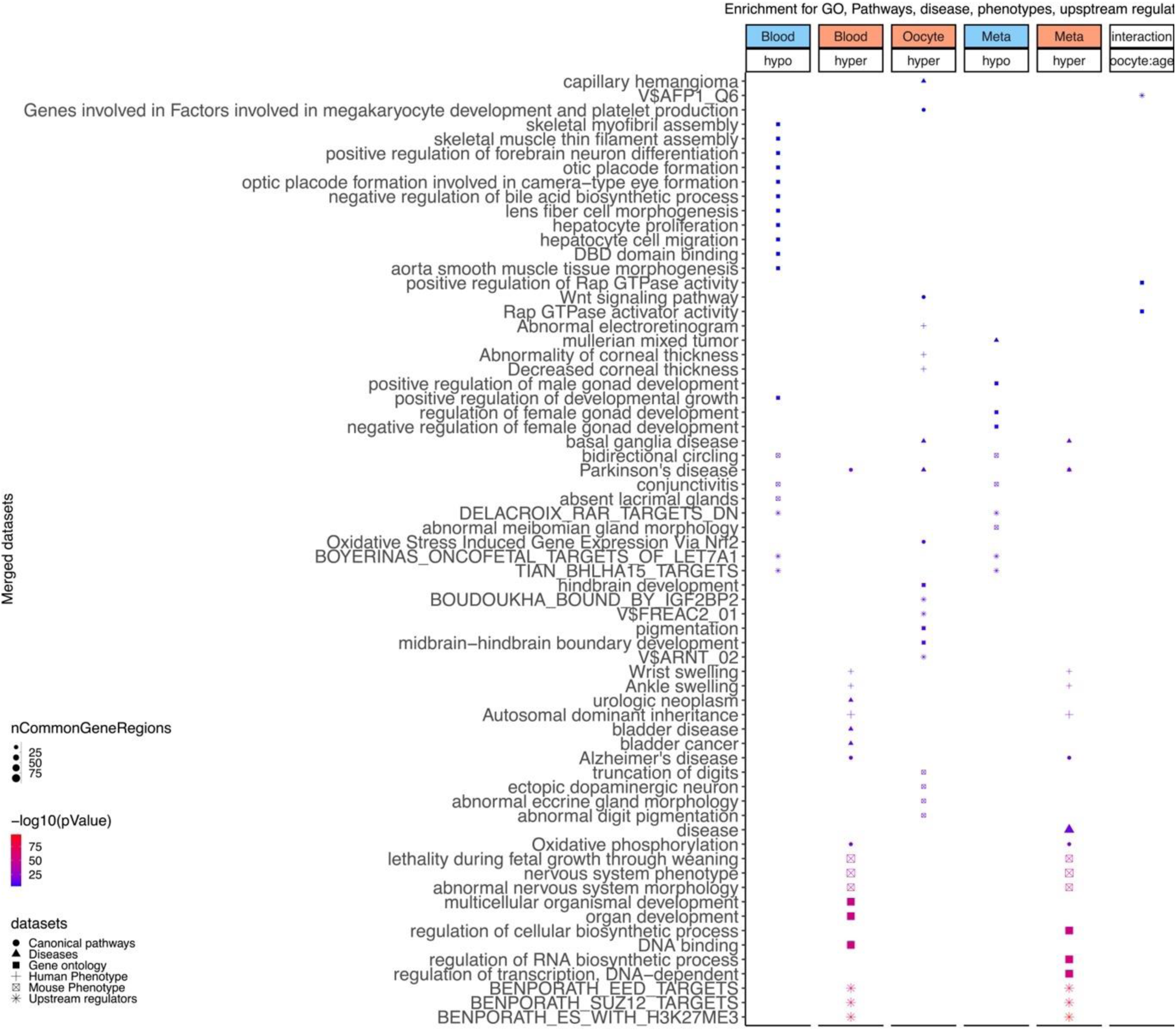
Enrichment analysis of the top aging-related CpGs in cattle. The analysis was done using genomic region of enrichment annotation tool [24]. The gene level enrichment was done using GREAT analysis [24] and human Hg19 background. The background probes were limited to 23775 probes that were mapped to the same gene in the cattle genome. The top 3 enriched datasets from each category (Canonical pathways, diseases, gene ontology, human and mouse phenotypes, and upstream regulators) were selected and further filtered for significance at p < 10^−4^.

**Supplementary Figure 3.**
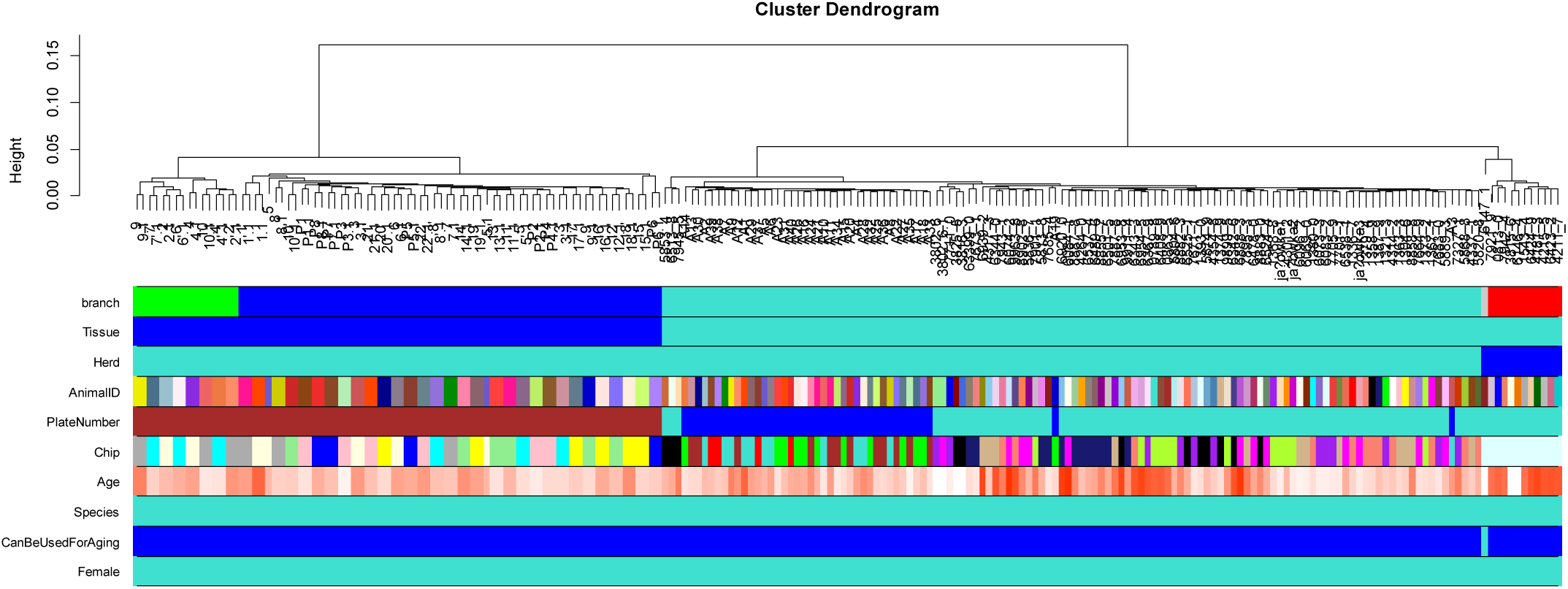
Unsupervised hierarchical clustering tree of dairy cattle. The comparison of the first two color bands underneath the tree reveals that the arrays (DNA samples) largely cluster by sample type. Tissue=turquoise corresponds to blood, Tissue=blue corresponds to oocytes. Weak evidence for sub clustering: the red cluster (rightmost cluster) can be explained: these samples come from cattle herd 2 (see the 3rd color band).

**Supplementary Figure 4.**
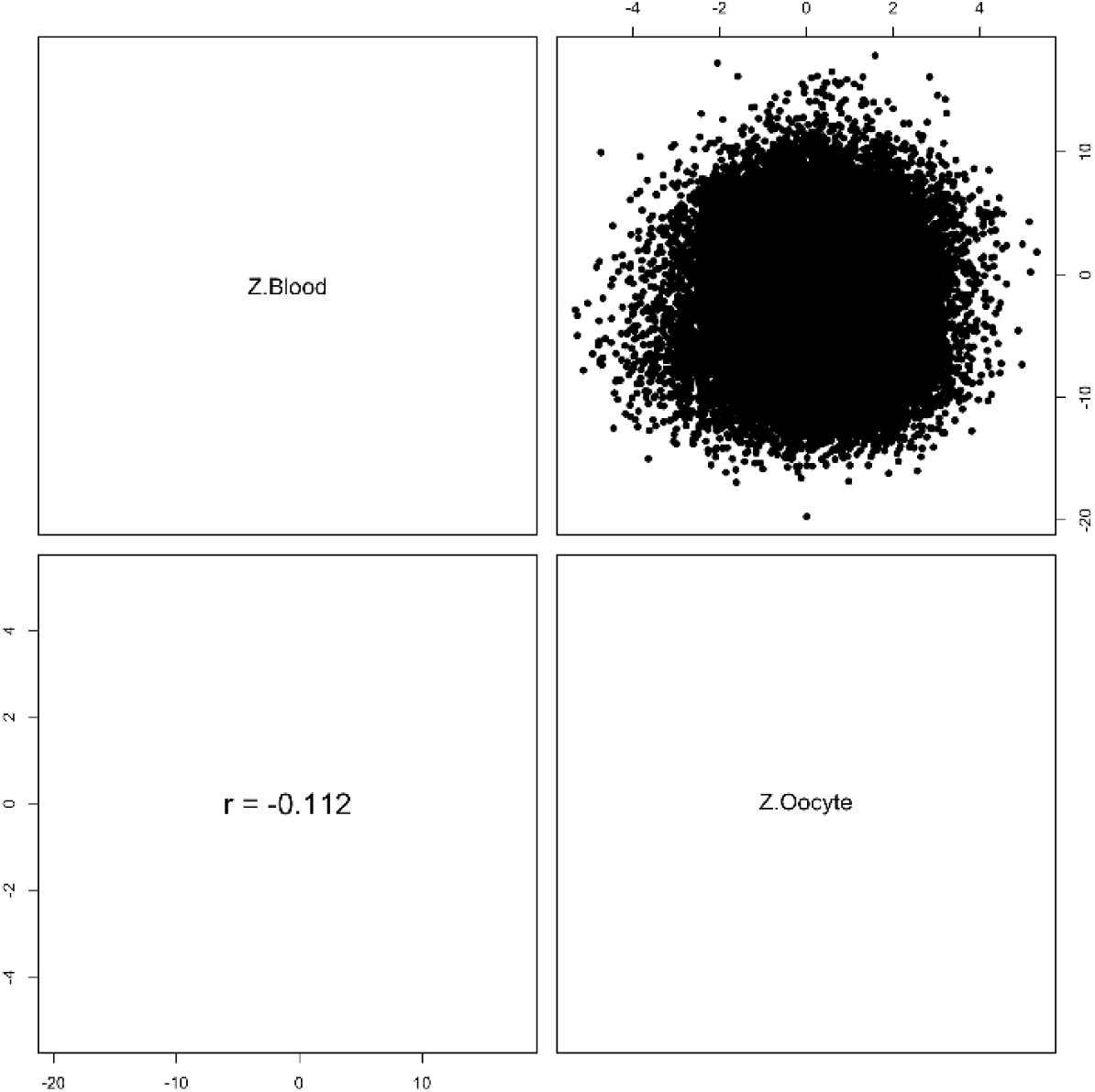
Correlating epigenome wide association study of age in oocytes and blood. Each dot corresponds to a CpG. Z statistics for a correlation test of age in cattle oocytes and blood.

**Supplementary Figure 5.**
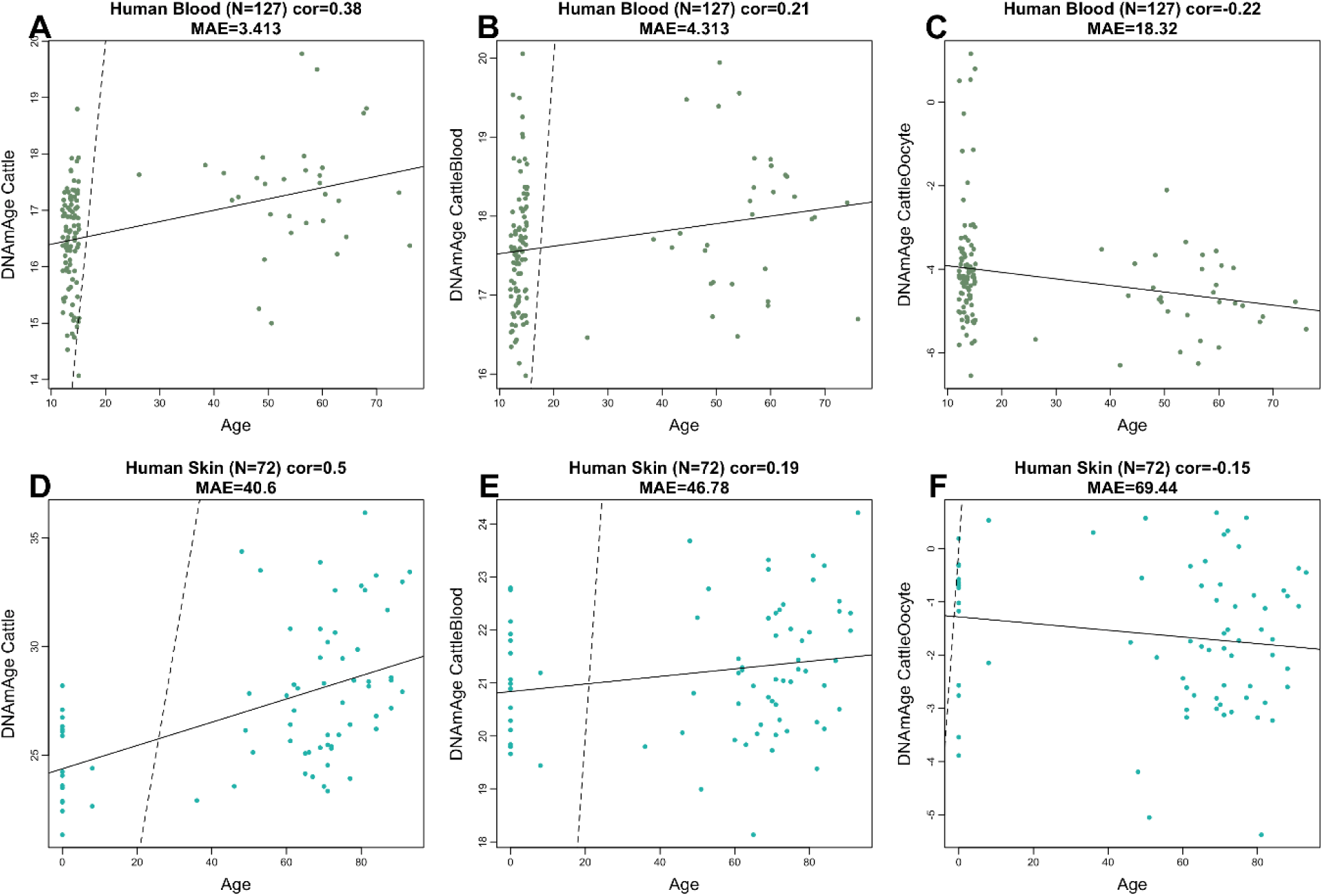
Pure cattle clocks applied to human samples. Each dot corresponds to a human tissue sample. A-C) Human blood samples. D-F) Human skin samples. The x axis reports the human chronological age. The y-axis corresponds to different epigenetic age estimates based on different cattle clocks. A,D) Multi-tissue clock for cattle. B,E) Blood clock for cattle. C,F) Oocyte clock for cattle.

**Supplementary Figure 6.**
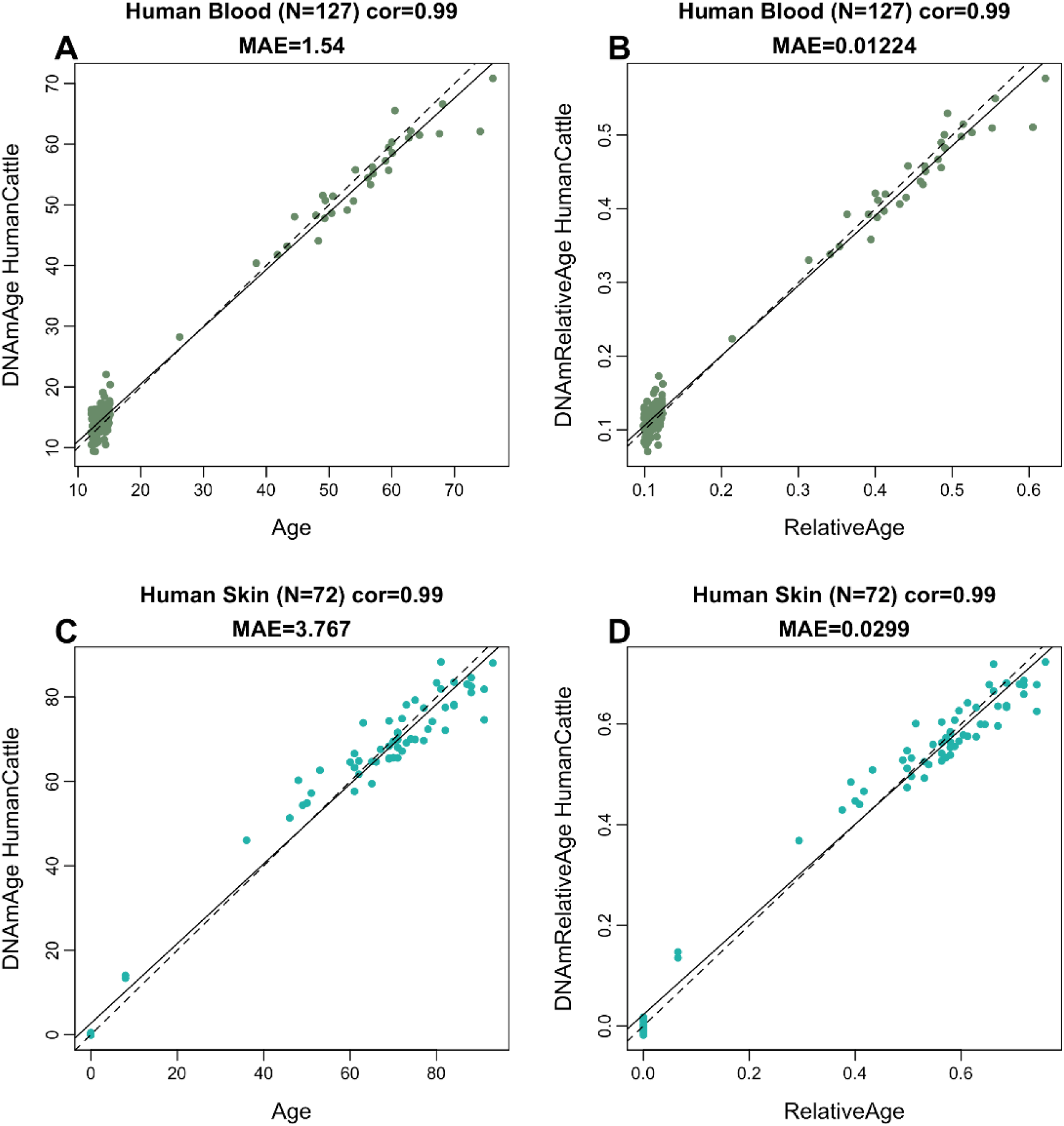
Human-cattle clocks applied to human samples. Each dot corresponds to a human tissue sample. Leave-one-human-sample-out (LOHO) cross validation was used to estimate age (y-axis). A,B) Human blood. C,D) Human skin samples. A,C) Human cattle clocks for chronological age. B,D) Human cattle clocks for relative age. The figure differs from Figure 1 in the following regards. First, it focuses on HUMAN samples as opposed to cattle samples which are presented in Figure 1E,H. Second, it focuses on a *subset* of human tissues (blood and skin) while Figure 1D,F used all human tissues (including liver, heart).

**Supplementary Figure 7.**
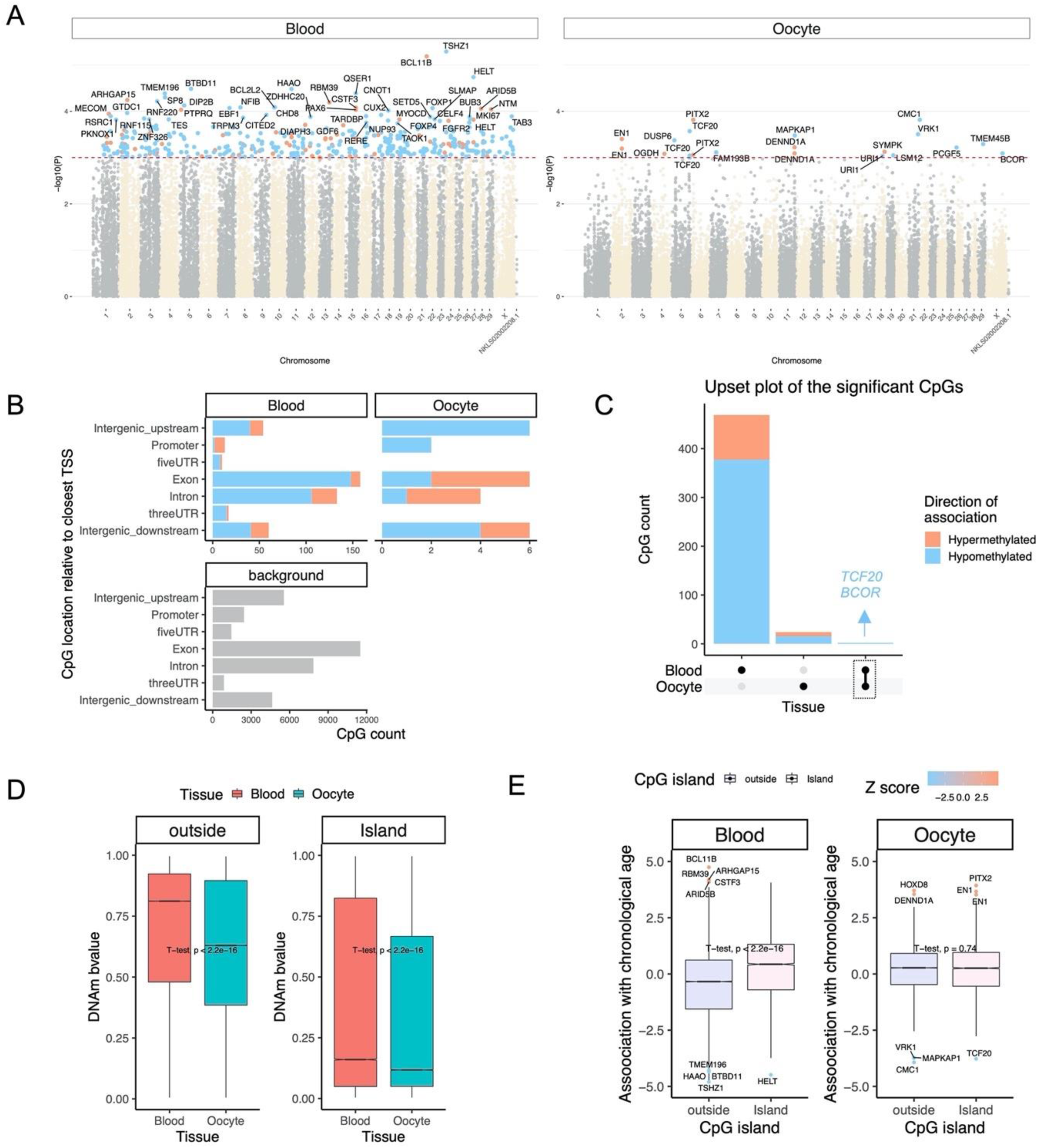
Age-dependent DNAm changes in blood and oocyte of the same animals. This analysis was performed on the subset (N=40) of the individuals who had both oocyte and blood samples. The observed pattern was similar to the observation in full population. A) Manhattan plots of the EWAS of chronological age. The coordinates are estimated based on the alignment of Mammalian array probes to Bos_taurus.ARS-UCD1.2 genome assembly. The red dotted line corresponds to a significance threshold of p < 10^−3^. Individual CpGs are colored in red or blue if they gain or lose methylation with age. The 50 most significant CpGs are labeled by neighboring genes. B) Location of top CpGs in each tissue relative to the closest transcriptional start site. Top CpGs were selected at p < 10^−3^. The number of selected CpGs: blood, 471; oocyte, 26. The grey color in the last panel represents the location of 34331 mammalian BeadChip array probes mapped to Bos_taurus.ARS-UCD1.2 genome. C) Upset plot representing the overlap of aging-associated CpGs based on meta-analysis or individual tissues. Neighboring genes of the overlapping CpGs were labeled in the figure. D) Oocytes have a general lower DNAm levels in both inside and outside of CpG islands than blood. The mean difference was examined by t-test. E) CpG islands have higher positive association with age (Z statistics of a correlation test for age) than CpGs located outside of the islands in blood but not in oocytes.

**Supplementary Figure 8.**
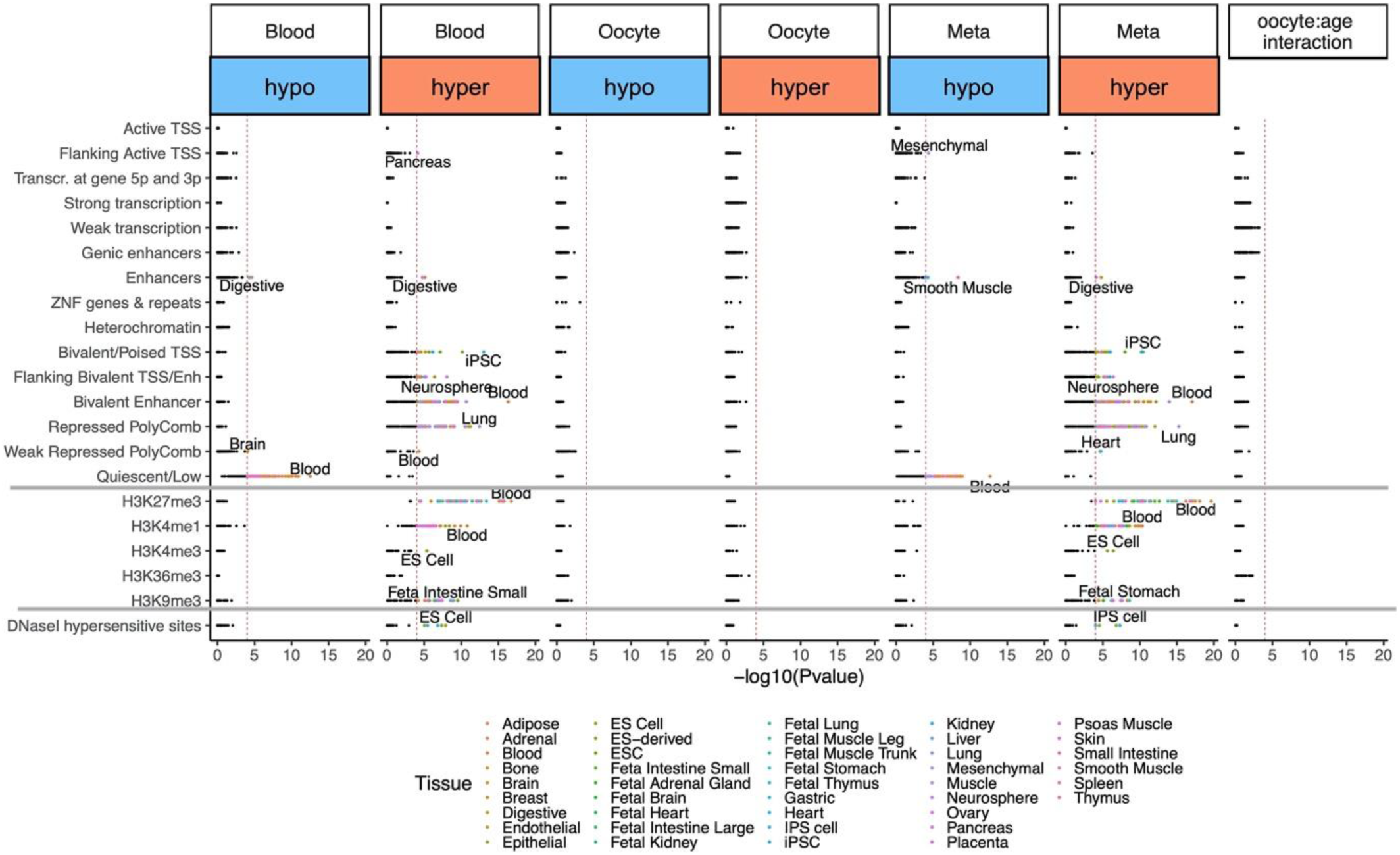
Enrichment of the cell type-specific chromatin states (15 states), histone 3 marks, and DNaseI I hypersensitivity sites for age-associated CpGs in cattles. Highlighted points indicates p < 10^−4^. Top tissue type for each significant mark are labeled. The analysis was performed with eForge V2.0 using the cattle genome as background.

**Supplementary Figure 9.**
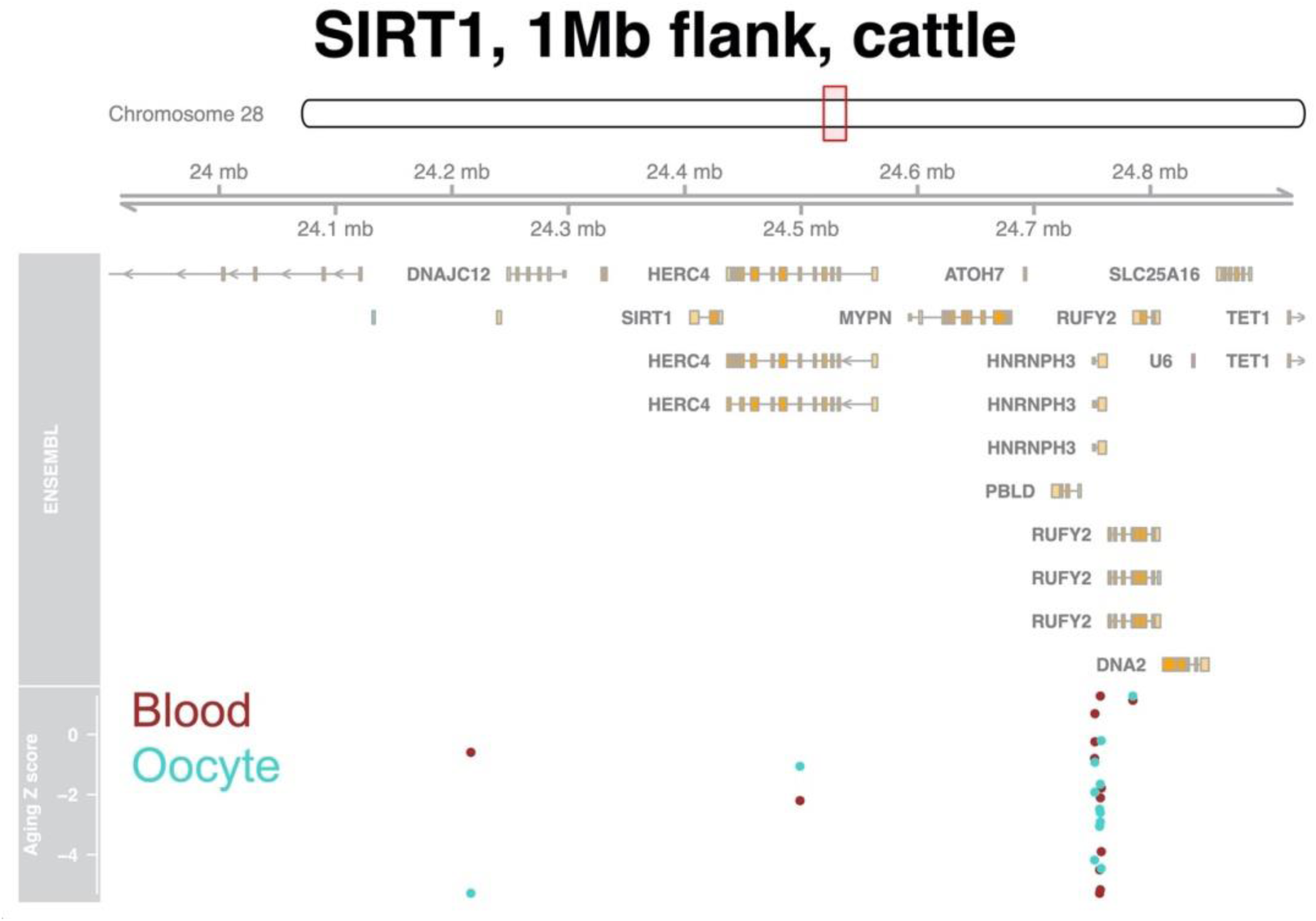
DNAm aging of the blood and oocyte in the SIRT1 flanking region of cattle. Given the absence of the SIRT1 gene on the mammalian array probes within a 1Mb flanking region of this gene were investigated. Several probes downstream of SIRT1 were hypomethylated with age in both blood and oocyte. One probe upstream of SIRT1 is only hypomethylated with age in the oocyte.

## Online content

Any methods, additional references, Nature Research reporting summaries, source data, extended data, supplementary information, acknowledgements, peer review information; details of author contributions and competing interests; and statements of data and code availability are available at:

## Methods

### Ethical authorization and animals

All animal procedures were carried out in accordance with the relevant guidelines at each institution. Specifically, procedures related to sample collection in Poland followed the EU Directive of the European Parliament and the Council on the protection of animals used for scientific purposes (22 September 2010; No 2010/63/EU), Polish Parliament Act on Animal Protection (21 August 1997, Dz.U. 1997 nr 111 poz. 724) with further novelization - Polish Parliament Act on the protection of animals used for scientific or educational purposes (15 January 2015, Dz.U. 2015 poz. 266). Blood and oocyte collection were approved by the Local Ethics Committee for Experiments on Animals, University of Warmia and Mazury in Olsztyn, Poland (Agreement No. LKE.065.27.2019). For animal procedures in the USA, approval from the University of Nebraska Institutional Animal Care and Use Committee was obtained (approval number is 1560). Samples were obtained from 357 female cattle (Bos Taurus). The animals were housed on the Dairy Farm of the Institute of Animal Reproduction and Food Research of Polish Academy of Sciences, (Wielki Las, Poland) and at the Eastern Nebraska Research and Extension Center at the University of Nebraska-Lincoln (Nebraska, USA). In the present study, Polish Red cattle was used for oocyte and blood collection. Further blood samples were also collected from the herd in Eastern Nebraska. In this herd, black Angus or composites of varying percentages of Simmental x Angus (black) or Red Angus where used as blood donors. Animals were free of Bovine Herpesvirus Type 1, Bovine Viral Diarrhea/Mucosal Disease, tuberculosis and Enzootic bovine leucosis.

### Blood collection and further processing

The blood samples from both herds were collected during routine animal management activities, either during the routine blood collection for disease prevention tests or during pregnancy diagnosis. Blood was collected into 8ml PAXgene Blood DNA Tubes (Quiagen, Cat No. 761115) and stored at −80°C until the shipment (USA Veterinary Permission Nr 138809) to the UCLA Technology Center for Genomics & Bioinformatics (Los Angeles, USA) for further analyses.

### Oocyte Collection and further processing

In total 80 Bovine ovaries were collected immediately *post mortem* from 40 cows which were selected for routine culling due to management reasons. Before the isolation of both ovaries from each cow, blood samples were also collected into 8ml PAXgene Blood DNA Tubes (Quiagen, Cat No. 761115) to generate both sample types from one donor. Isolated ovaries were kept on ice and immediately transported to the laboratory. Afterwards, immature bovine cumulus–oocyte complexes (COCs) were recovered by aspirating ovarian follicles in the diameter of 2–8 mm. Cumulus cells were removed from COCs by pipetting them for 5min in a Petri dish containing 500 μl of Phosphate Buffered Saline (PBS) with 0.1% hyaluronidase (Sigma-Aldrich, St. Louis, MO, USA). Denuded oocytes with clean *zona pellucida* were then processed for genomic DNA isolation.

### Genomic DNA isolation

Genomic DNA was isolated from blood and oocyte samples using the DNeasy Blood and Tissue Kit (Qiagen, Cat No. 69506) and subsequently bisulfite converted using the EZ DNAMethylation Kit (ZymoResearch, Irvine, CA, USA). Bisulfite-treated samples were processed using the custom array.

### DNA methylation data

All DNAm data used was generated using the custom Illumina chip “HorvathMammalMethylChip40”, also known as mammalian methylation array. The mammalian methylation array is an attractive tool for DNAm assessment in primates, because it comprises 38K probes, including nearly ∼36 K probes targeting CpG sites in highly conserved regions in mammals. Briefly, two thousands out of 38 K probes were selected based on their utility for human biomarker studies: these CpGs, which were previously implemented in human Illumina Infinium arrays (EPIC, 450K) were selected due to their relevance for estimating age, blood cell counts, or the proportion of neurons in brain tissue. The remaining 35,988 probes were chosen to assess cytosine DNA methylation levels in mammalian species. Highly conserved CpGs across 50 mammalian species were selected: 33,493 Infinium II probes and 2,496 Infinium I probes. Each probe of these highly conserved probes was designed to cover a certain subset of species, such that overall all species have a high number of probes (A. Arneson, J. Ernst, Horvath, in preparation). The particular subset of species for each probe is provided in the chip manifest file can be found at Gene Expression Omnibus (GEO) at NCBI as platform GPL28271. The SeSaMe normalization method was used to define beta values for each probe (Zhou et al. 2018).

### Probe mapping and annotation

The probe sequences were aligned to cattle (*Bos taurus*) ARS-UCD1.2 genome using QUASR package [25] with the assumption for bisulfite conversion treatment of the genomic DNA. The QUASR (a wrapper for Bowtie2) was run with parameters -k 2 --strata --best -v 3 and bisulfite = “undir” to align the probe sequences to each prepared genome. Following the alignment, the CpGs were annotated based on the distance to the closest transcriptional start site using the Chipseeker package [26]. A gff file with these was created using these positions, sorted by scaffold and position, and compared to the location of each probe in BAM format. Genomic location of each CpG was categorized as intergenic, 3’ UTR, 5’ UTR, promoter region (minus 10 kb to plus 1000 bp from the nearest TSS), exon, or intron.

### Epigenome-wide association studies of age

The DNAm changes were examined for association with age using the R function “standardScreeningNumericTrait” from the “WGCNA” R package [27] in each tissue. The results were combined using Stouffer’s meta-analysis method.

Downstream enrichment analysis of the significant differentially methylated positions (DMP) was performed for transcriptional factor (TF) motifs, and merged datasets of gene ontology, canonical pathways, upstream regulators, diseases, and phenotypes using the neighboring gene sets. For TF enrichment, we used MEME motif discovery algorithm to identify the background motifs and included the CpGs with FIMO (Find Individual Motif Occurrence) p-value < 10^−5^. The overlap of probe sets and transcriptional factor motifs were tested using a hypergeometric test (phyper).

The gene-level enrichment was done using GREAT analysis [24] and human Hg19 background, but prefiltered to 23775 probes that were mapped to the same gene in both human and cattle genomes.

## References

1 te Velde ER, Pearson PL. The variability of female reproductive ageing. Human Reproduction Update, 8, 141–154. doi:10.1093/humupd/8.2.141; PMID:12099629

2 Selesniemi K, Lee HJ, Tilly JL. Moderate caloric restriction initiated in rodents during adulthood sustains function of the female reproductive axis into advanced chronological age. Aging Cell. 2008;7:622–629. doi: 10.1111/j.1474-9726.2008.00409; PMID:18549458

3 Garg,N, Sinclair DA. Oogonial stem cells as a model to study age-associated infertility in women. Reproduction, fertility, and development 2015. doi:10.1071/RD144614; PMID: 25897831

4 Baired DT, Collins J, Egozcue J, Evers LH, Gianaroli L, Leridon H, Sunde A, Templeton A, Van Steirteghem A, Cohen J, Crosignani PG, Devroey P, Diedrich K, Fauser BC, Fraser L, Glasier A, Liebaers I, Mautone G, Penney G, Tarlatzis B. Fertility and ageing., Hum Reprod Update. 2005 May-Jun;11(3):261–76. doi: 10.1093/humupd/dmi006; PMID:15831503

5 Navot D, Fox JH, Williams M, Brodman M, Friedman F Jr, Cohen CJ. The concept of uterine preservation with ovarian malignancies. Lancet. 1991 Jun 8;337(8754):1375–7. doi: 10.1016/0140-6736(91)93060-m; PMID:1870826

6 Dodson MG,Minhas BS, Curtis SK, Palmer TV, Robertson JL. Spontaneous zona reaction in the mouse as a limiting factor for the time in which an oocyte may be fertilized. J. In Vitro Fert Embryo Transf. 1989;6:101–106; PMID: 2723502

7 Xu Z, Abbott A, Kopf GS, Schultz RM, Ducibella T. Spontaneous activation of ovulated mouse eggs: time-dependent effects on M-phase exit, cortical granule exocytosis, maternal messenger ribonucleic acid recruitment, and inositol 1,4,5-trisphosphate sensitivity.Biol Reprod. 1997;57:743–750. doi.org/10.1095/biolreprod57.4.743; PMID: 9314575

8 Kikuchi K, Naito K, Noguchi J, Shimada A, Kaneko H, Yamashita M, Aoki F, Tojo H, Toyoda Y. Maturation/M-phase promoting factor: a regulator of aging in porcine oocytes. Biol Reprod. 2000;63:715–722. doi.org/10.1095/biolreprod63.3.715; PMID:10952912

9 Liu L, Blasco MA, Keefe DL. Requirement of functional telomeres for metaphase chromosome alignments and integrity of meiotic spindles., EMBO Reports (2002)3:230–234. doi.org/10.1093/embo-reports/kvf055; PMID:11882542

10 Pellestor F, Andréo B, Arnal F, Humeau C, Demaille J. Maternal aging and chromosomal abnormalities: new data drawn from in vitro unfertilized human oocytes. Hum Genet. 2003;112:195–203. doi.10.1007/s00439-002-0852-x; PMID:12522562

11 López-Otín C, Blasco MA, Partridge L, Serrano M, Kroemer G. The hallmarks of aging. Cell. 2013;153(6):1194–1217. doi:10.1016/j.cell.2013.05.039; PMID: 23746838

12 Petro EM, Covaci A, Leroy JL, Dirtu AC, De Coen W, and Bols PE. Occurrence of endocrine disrupting compounds in tissues and body fluids of Belgian dairy cows and its implications for the use of the cow as a model to study endocrine disruption. Sci Total Environ 2010: 408; 5423–5428. DOI: 10.1016/j.scitotenv.2010.07.051. PMID: 20709361

13 Adams GP, Singh J, and Baerwald AR. Large animal models for the study of ovarian follicular dynamics in women. Theriogenology 2012: 78; 1733–1748. DOI: 10.1016/j.theriogenology.2012.04.010. PMID: 22626769

14 Marshall KL, Rivera RM. The effects of superovulation and reproductive aging on the epigenome of the oocyte and embryo. Mol Reprod Dev. 2018;85(2):90–105. doi:10.1002/mrd.22951

15 Horvath S. DNA methylation age of human tissues and cell types. Genome Biol 14, R115 (2013). doi: 10.1186/gb-2013-14-10-r115; PMID: 24138928

16 Petkovich DA, Podolskiy DI, Lobanov AV, Lee SG, Miller RA, Gladyshev VN. Using DNA Methylation Profiling to Evaluate Biological Age and Longevity Interventions. Cell Metab. 2017;25(4):954–960.e6. doi:10.1016/j.cmet.2017.03.016; PMID: 28380383

17 Goodarzi MO, Jones MR, Li X, Chua AK, Garcia OA, Chen YD, Krauss RM, Rotter JI, Ankener W, Legro RS, Azziz R, Strauss JF 3rd, Dunaif A, Urbanek M. Replication of association of DENND1A and THADA variants with polycystic ovary syndrome in European cohorts. J Med Genet. 2012 Feb;49(2):90–5. doi: 10.1136/jmedgenet-2011-100427. PMID: 22180642;

18 Eriksen MB, Nielsen MF, Brusgaard K, et al. Genetic alterations within the DENND1A gene in patients with polycystic ovary syndrome (PCOS). PLoS One. 2013;8(9):e77186. Published 2013 Sep 27. doi:10.1371/journal.pone.0077186. PMID: 24086769

19 Jaśkiewicz A, Pająk B, Orzechowski A. The Many Faces of Rap1 GTPase. Int J Mol Sci. 2018;19(10):2848. Published 2018 Sep 20. doi:10.3390/ijms19102848. PMID: 30241315.

20 Tapia A, Gangi LM, Zegers-Hochschild F, et al. Differences in the endometrial transcript profile during the receptive period between women who were refractory to implantation and those who achieved pregnancy. Hum Reprod. 2008;23(2):340–351. doi:10.1093/humrep/dem319. PMID: 18077318

21 Horvath, S. DNA methylation age of human tissues and cell types. Genome Biol 14, 3156 (2013). https://doi.org/10.1186/gb-2013-14-10-r115

22 Iljas JD, Wei Z, Homer HA. Sirt1 sustains female fertility by slowing age-related decline in oocyte quality required for post-fertilization embryo development [published online ahead of print, 2020 Jul 30]. Aging Cell. 2020;e13204. doi:10.1111/acel.13204

23 Fahy GM, Brooke RT, Watson JP, et al. Reversal of epigenetic aging and immune-senescent trends in humans. Aging Cell. 2019;18(6):e13028. doi:10.1111/acel.13028

24 McLean, C. Y., Bristor, D., Hiller, M., Clarke, S. L., Schaar, B. T., Lowe, C. B., Bejerano, G. (2010). GREAT improves functional interpretation of cis-regulatory regions. Nat Biotechnol, 28(5), 495–501. doi:10.1038/nbt.1630

25 Gaidatzis, D., Lerch, A., Hahne, F., & Stadler, M. B. (2015). QuasR: quantification and annotation of short reads in R. Bioinformatics, 31(7), 1130–1132. doi:10.1093/bioinformatics/btu781

26 Yu, G., Wang, L. G., & He, Q. Y. (2015). ChIPseeker: an R/Bioconductor package for ChIP peak annotation, comparison and visualization. Bioinformatics, 31(14), 2382–2383. doi:10.1093/bioinformatics/btv145

27 Langfelder, P., & Horvath, S. (2008). WGCNA: an R package for weighted correlation network analysis. BMC Bioinformatics, 9(1), 559. Retrieved from http://www.biomedcentral.com/1471-2105/9/559

